# Prevalence of Zika virus infection in wild African primates

**DOI:** 10.1101/077628

**Authors:** Connor R. Buechler, Adam L. Bailey, Andrea M. Weiler, Gabrielle L. Barry, Anna J. Jasinska, Nelson B. Freimer, Cristian Apetrei, Jane E. Phillips-Conroy, Clifford J. Jolly, Jeffrey Rogers, Thomas C. Friedrich, David H. O’Connor

## Abstract

The recent spread of Zika virus (ZIKV) is alarming due to its association with birth defects. Though the natural reservoir of ZIKV remains poorly defined, the virus was first described in a captive “sentinel” macaque in Africa. Here, we examined blood from 239 wild African monkeys and found variable seropositivity.

## INTRODUCTION

Zika virus (ZIKV) is a positive-sense single-stranded RNA virus that recently emerged to cause widespread human infection in Micronesia and the Americas. However, the cause of ZIKV’s recent expansion is poorly understood and its natural reservoir remains poorly defined. ZIKV was first isolated from a captive rhesus macaque (*Macaca mulatta*) in Uganda in 1947 (1). Since then, ZIKV has been isolated from humans, monkeys, and mosquitoes living in Africa, Southeast Asia, Micronesia, and the Americas (2). Monkeys in Africa often live in close association with humans and are likely involved in the zoonotic transmission of flaviviruses, including ZIKV, mediated by mosquito vectors (3). While many serosurveys showed the presence of ZIKV in both humans and non-human primates in Africa in the 1950s-1980s (2), these surveys have not been updated since. In addition, serosurveys from this period were prone to significant cross-reactivity with other flaviviruses such as dengue virus (DENV), West Nile virus (WNV), and yellow fever virus (YFV), complicating their interpretation (2). We sought to better understand ZIKV prevalence in wild African primates using unbiased deep-sequencing, ZIKV-specific quantitative RT-PCR, and a ZIKV-specific antibody-capture assay, assessing for antibody cross-reactivity to other flaviviruses using serum from experimentally infected animals.

## STUDY METHODOLOGY

Four populations of wild non-human primates were included in this study -- African green monkeys from across South Africa (*Chlorocebus pygerythrus*) and The Gambia (*Chlorocebus sabaeus*), hybrid kinda × grayfooted-chacma baboons from Kafue National Park in Zambia (*Papio kindae* × *Papio ursinus griseipes*), and yellow baboons from Mikumi National Park in Tanzania (*Papio cynocephalus*). All sampling was performed according to regulations set by the Animal Welfare Act, and all collection methods were approved by appropriate Animal Care and Use Committees. Briefly, animals were individually trapped, sedated, and bled, resulting in a total of 239 plasma samples. Samples from Tanzania were collected in 1985 and 1986; samples from The Gambia, South Africa, and Zambia were collected between 2010 and 2014.

Samples from South Africa, Zambia, and Tanzania were processed for deep sequencing as described previously (4). Briefly, viral RNA was isolated from approximately 200 µl of plasma and cDNA synthesis was primed using random hexamer primers. Samples were fragmented, and sequencing adaptors were added using a Nextera DNA sample preparation kit (Illumina, San Diego, CA, USA). Deep sequencing was performed on an Illumina MiSeq. Sequence data were analyzed using CLC Genomics Workbench (version 6.5) software (CLC Bio, Aarhus, Denmark) and Geneious R9 software (Biomatters, Auckland, New Zealand). Low-quality (Phred quality score, <Q30) and short reads (<100 bp) were removed. Samples were then compared against all viral sequences in the NCBI GenBank database as of June 22, 2016 (5) using BLAST. ZIKV reads were not identified in any of the samples investigated by this method.

All samples were tested for ZIKV-specific immunoglobulin-G (IgG) against recombinant ZIKV nonstructural protein 1 (NS1) using ELISA kits designed by XPressBio (XPressBio, Frederick, MD, USA). Samples were diluted 1:50 and the assay was performed according to the manufacturer’s instructions, with all tests performed in duplicate. Specificity was independently verified by testing serum from animals experimentally infected with ZIKV (n=6), DENV-1 (n=6), DENV-2 (n=5), DENV-3 (n=12), DENV-4 (n=6); vaccinated for YFV (n=6); or raised in captivity with no known exposure to flaviviruses (n=12). Sera from animals vaccinated with YFV showed no cross-reactivity with this antibody detection assay. However, cross-reactivity was noted with all DENV serotypes except DENV-2 (supplemental material). False-positive reactions from animals previously infected with DENV showed an average change in optical density (ΔOD) between positive and negative wells of 0.815 with a standard deviation of 0.345, which was above the manufacturer-recommended positivity threshold of ΔOD > 0.3. We adopted a new experimentally-determined cutoff at the upper limit of the 95% confidence interval of this population (ΔOD = 1.49) to more accurately distinguish ZIKV exposure from DENV exposure. All samples from animals experimentally infected with ZIKV had ΔOD values that were greater than this limit, making the ELISA test 100% sensitive and 100% specific based on experimentally infected animals with known clinical histories. Amongst the wild monkeys tested, the highest seroprevalence was seen in the Gambian vervets, where 4 out of 25 animals (16%) showed evidence of prior exposure to ZIKV. Two of the 41 Tanzanian yellow baboons tested (4.9%) were also ZIKV seropositive. None of the samples from Zambian baboons (n=14) or South African AGMs (n=159) were positive for antibodies to ZIKV.

Samples from The Gambia and any from South Africa that tested above the manufacturer-recommended threshold of ΔOD > 0.3 for anti-ZIKV IgG by ELISA were also subjected to highly-sensitive quantitative RT-PCR, as described previously (6), using primers and probe as follows: Forward – CGYTGCCCAACACAAGG; Reverse – CCACYAAYGTTCTTTTGCABACAT; Probe-AGCCTACCTTGAYAAGCARTCAGACACYCAA (Aliota et al. *under review*). The qRT-PCR was performed using the SuperScript III Platinum one-step quantitative RT–PCR system (Invitrogen, Carlsbad, CA, USA) on the LightCycler 480 instrument (Roche Diagnostics, Indianapolis, IN, USA). Virus concentration was determined by interpolation onto an internal standard curve composed of seven 10-fold serial dilutions of a synthetic ZIKV RNA fragment based on ZIKV strain H/PF/2013. ZIKV RNA was not detected in any of the samples tested.

## CONCLUSION

We assessed the prevalence of ZIKV infection in four wild primate populations in Africa: baboons (genus *Papio*) from Tanzania and Zambia and AGMs (genus *Chlorocebus*) from South Africa and The Gambia. We found evidence for prior ZIKV infection in Tanzanian yellow baboons and Gambian AGMs, but not in the other primate populations. These results confirm that ZIKV infection in wild primates is not constrained by particular host species but instead may be influenced by other factors such as geography, as we found antibodies to ZIKV to be more common in primates living in the northern parts of sub-Saharan Africa. The geographical range of ZIKV-transmitting mosquitoes may provide an explanation for this observation. Common ZIKV vectors such as *Aedes aegypti* and *Aedes albopictus* are less common in both Zambia and South Africa compared to Tanzania and The Gambia (7), and the range of *Aedes africanus* is largely limited to tropical forests in equatorial Africa (8).

As ZIKV causes an acute viral infection, it is not surprising that our cross-sectional study did not detect active infection by deep sequencing or qRT-PCR. That we did not find evidence of active ZIKV infection in antibody-positive animals is consistent with recent data showing that ZIKV-specific antibodies are sufficient to protect against infection, such that antibody and viral nucleic acids would not likely coexist (9).

Extensive cross-reactivity among anti-flaviviral antibodies has long presented a significant barrier to accurate determination of the seroprevalence of ZIKV. Thus, the increased specificity of the ELISA with the cutoff value developed here represents an important advancement for this field. Indeed, the ZIKV seroprevalence estimates reported here are well below many of those noted in previous studies (2), possibly reflecting the lower false-positive rate of this method and providing a more accurate sense of the prevalence of ZIKV in the communities surveyed. This method can be used for larger serosurveys of wild animals to more accurately determine the prevalence of ZIKV infection. Such studies will be vital for further understanding the natural reservoir and transmission dynamics of ZIKV in Africa and elsewhere.

**Figure 1.**
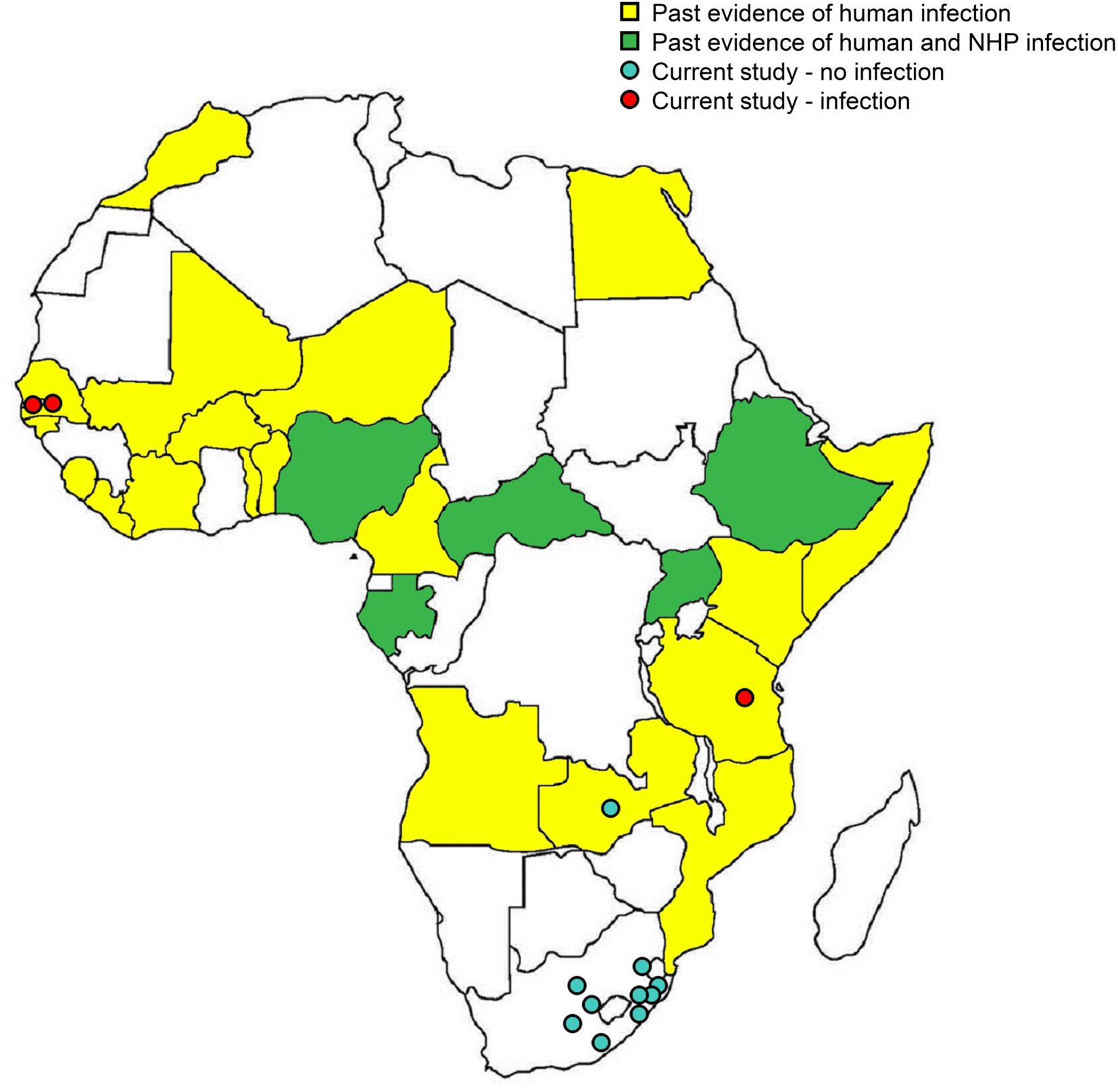
ZIKV Detection in African Primates. Countries where human ZIKV infection has been detected are shown in yellow; countries where human and non-human primate ZIKV infection has been detected are shown in green; locations in which non-human primate ZIKV infection was detected in the current study are shown with red dots; locations in which non-human primate ZIKV infection was not detected in the current study are shown with blue dots. Past studies were reviewed in Musso & Gubler, 2016 (2).

## Acknowledgments

We would like to acknowledge the advice and support of the entire Zika Experimental Science Team. More information can be found at zika.labkey.com. We thank the DHHS/PHS/NIH (R01Al116382-01A1 to D.H.O’C) for funding. We also thank the P51OD011106 awarded to the WNPRC, Madison-Wisconsin. This research was conducted in part at a facility constructed with support from the Research Facilities Improvement Program (grant numbers RR15459-01 and RR020141-01). The collection of blood samples from Mikumi baboons was funded by NSF grant BNS83-03506 to J.P.-C. (Washington University). Vervet samples were collected through the UCLA Systems Biology Sample Repository funded by NIH grants R01RR016300 and R01OD010980 to N.B.F. We thank the Gambia Department of Parks & Wildlife Management, Ministry of Forestry & the Environment, Department of Environmental Affairs, South Africa; Department of Economic Development and Environmental Affairs, Eastern Cape; Department of Tourism, Environmental and Economic Affairs, Free State Province; the Ezemvelo KZN Wildlife in KwaZulu Natal Province; and the Department of Economic Development, Environment and Tourism, Limpopo Province. We also thank the Zambian Wildlife Authority for permission to conduct research in Kafue National Park and the Tanzanian National Parks Authority (TANAPA) for permission to work at Mikumi National Park.

## Author Bio

Connor Buechler is an Associate Research Specialist at the University of Wisconsin School of Medicine and Public Health with a primary research interest in infectious disease surveillance and prevention.

